# Development of a growth coupled dynamic regulation network balancing malonyl-CoA node to enhance (2*S*)-naringenin synthesis in *E. coli*

**DOI:** 10.1101/2020.07.07.192633

**Authors:** Shenghu Zhou, Shuo-Fu Yuan, Priya H Nair, Hal S Alper, Yu Deng, Jingwen Zhou

## Abstract

Generally, high- and low-performance nongenetic variants and young and aged cells co-existed in culture at all growth phases. In this regard, individually and dynamically regulating the metabolic flux of single cells based on their cellular state is highly useful for improving the performance of populations. However, balancing the trade-offs between biomass formation and compound over-production requires sophisticated dynamic regulation networks (DRNs) which can be challenging. Here, we developed a growth coupled NCOMB (Naringenin-Coumaric acid-Malonyl-CoA-Balanced) DRN with systematic optimization of (2*S*)-naringenin and *p*-coumaric acid-responsive regulation pathways for real-time control of intracellular supply of malonyl-CoA. In doing so, the acyl carrier protein was used as a new critical node for fine-tuning malonyl-CoA consumption instead of fatty acid synthase. Following directed evolution of the NCOMB DRN, we obtained a strain with cumulative 8.4-fold improvement in (2*S*)-naringenin production with balanced cell growth, demonstrating the high efficiency of this system for improving pathway production.

## 1 Introduction

Microbial production of value-added biochemicals often requires the development of a robust, efficient, and high-performance strain. To this end, static metabolic engineering strategies including multi-module optimization (*1*), promoter and RBS engineering (*2*), and CRISPR/Cas9-based genome engineering (*3*) are commonly employed for the rewiring of metabolic flux to maximize production. Despite significant successes in the literature and industry using these static strain engineering paradigms, it remains difficult to balance the trade-offs between growth and production. To address this balance within static strain engineering approaches, researchers often utilize culture-level changes including dynamic environmental changes such as incubation temperature, osmolarity, and nutrient availability or oscillating levels of intermediate metabolites (*4*). As an alternative, genetic approach, the use of transcription factor-based biosensors to monitor cell growth and/or metabolism provides the potential to enable a responsive mode to metabolism termed dynamic metabolic regulation (*5*).

Using the principles of dynamic regulation, a series of control systems (typically enabled by biosensors) fine-tune the cells to synthesize a target compound at maximum production levels (*4*). These control systems can be dynamically regulated at either the environmental or metabolic level. Generally, biosensors that could respond to universal, environmental induction factors such as temperature (*6*), dissolved oxygen (*5*), and light (*7*) enable construction of environment-responding dynamic regulation networks (DRNs). For example, Zhao *et al.* established a blue light inducible transcription factor (VP16-EL222) controlled OptoEXP system for isobutanol production and demonstrated a 4-fold improvement in the titer compared with the strain lacking dynamic regulatory circuits (*8*). Likewise, temperature is a common environment inducer. Using the promoter P_R_/P_L_ system along with the repressor protein CI857, temperature was able to dynamically control the expression of isocitrate dehydrogenase (*icd*) thus achieving a balance between itaconic acid biosynthesis and cell growth (*9*). The cell factories constructed in these studies respond to environment changes, however, environmental signals represent the integration of cellular state and thus not explicitly reflect the differences between intracellular states.

More specifically, fermentation conductions such as cell density represent the overall state of the cell population. In reality, a culture contains a mixture of high- and low-performance nongenetic variants and young and aged cells in all growth phases (*10*). As a result, precisely regulating metabolic flux based on the intracellular state can help improve overall productivity within a fermentation. To this end, it is important to identify biosensors that can directly monitor the intracellular metabolic state of single cells and thus individually regulate gene expression within single cells in response to a critical metabolite level (*4*). Recent examples of this strategy have been used to balance malonyl-CoA as it serves as a central metabolic precursor. For example, by coupling the glucose-responsive promoter (P_HXT1_) with a malonyl-CoA sensitive transcriptional repressor (FapR), David *et al.* constructed a DRN to drive the flux from the malonyl-CoA pool toward 3-hydroxypropionic acid formation, leading to a 10-fold increase in titer (*11*). Additionally, Xu *et al.* developed an oscillator to dynamically balance malonyl-CoA pool through controlling the expression of acetyl-CoA carboxylase (ACC) and fatty acid synthase (*12*). In their report, the best producer displayed a 2.1-fold improvement in fatty acids titer (3.86 g/L) (*13*). While successful in improving product titers, metabolite-responsive DRNs only monitor the fluctuating level of a metabolite without coupling with the growth profile of single cells, usually at the expense of growth.

In contrast to these approaches, the work present here demonstrates a dynamic optimization coupling that utilizes a growth coupled single cell sensing and regulating DRNs cascade to improve the production of the flavonoid (2*S*)-naringenin. Specifically, this DRN cascade focuses on malonyl-CoA, a crucial metabolite used as a carbon-chain elongation unit for important compounds such as flavonoids (*14*), antibiotics (*15*), and fatty acids (*12*). However, the availability of intracellular malonyl-CoA is limited and reduces from 0.23 to 0.01 nmol/(mg dry wt) when cells transit from exponential to stationary stage (*16*). Traditional approaches to increase malonyl-CoA content usually represses the fatty acid biosynthetic pathway (*17*). However, excessive repression of this pathway leads to impaired cell growth. Additionally, oversupply of malonyl-CoA causes an undesired malonylation of the proteome, thus establishing a further carbon burden in engineered *E. coli*. (*18*). As a result, malonyl-CoA must be in an exquisite balance within cells. In this study, we designed and optimized a metabolite-responsive NCOMB (Naringenin-Coumaric acid-Malonyl-CoA-Balanced) DRN which can real-time monitor the concentration of (2*S*)-naringenin and *p*-coumaric acid to dynamically enhance the production of malonyl-CoA at all growth phases. The designed DRN includes three modules: (1) module A that functionally produces (2*S*)-naringenin, (2) module B that dynamically amplifies the metabolic flux toward (2*S*)-naringenin synthetic pathway with coupling cell growth, and (3) module C that dynamically responds to *p*-coumaric acid and enhances the production of malonyl-CoA during late stages of fermentation (**Fig. 1**). In doing so, we successfully demonstrate the ability of NCOMB DRN to achieve the single cell dynamic regulation and achieved a balance of (2*S*)-naringenin production as well as cell growth based on the level of intracellular metabolite, which could further facilitate the advancement of DRN design.

**Figure 1.**
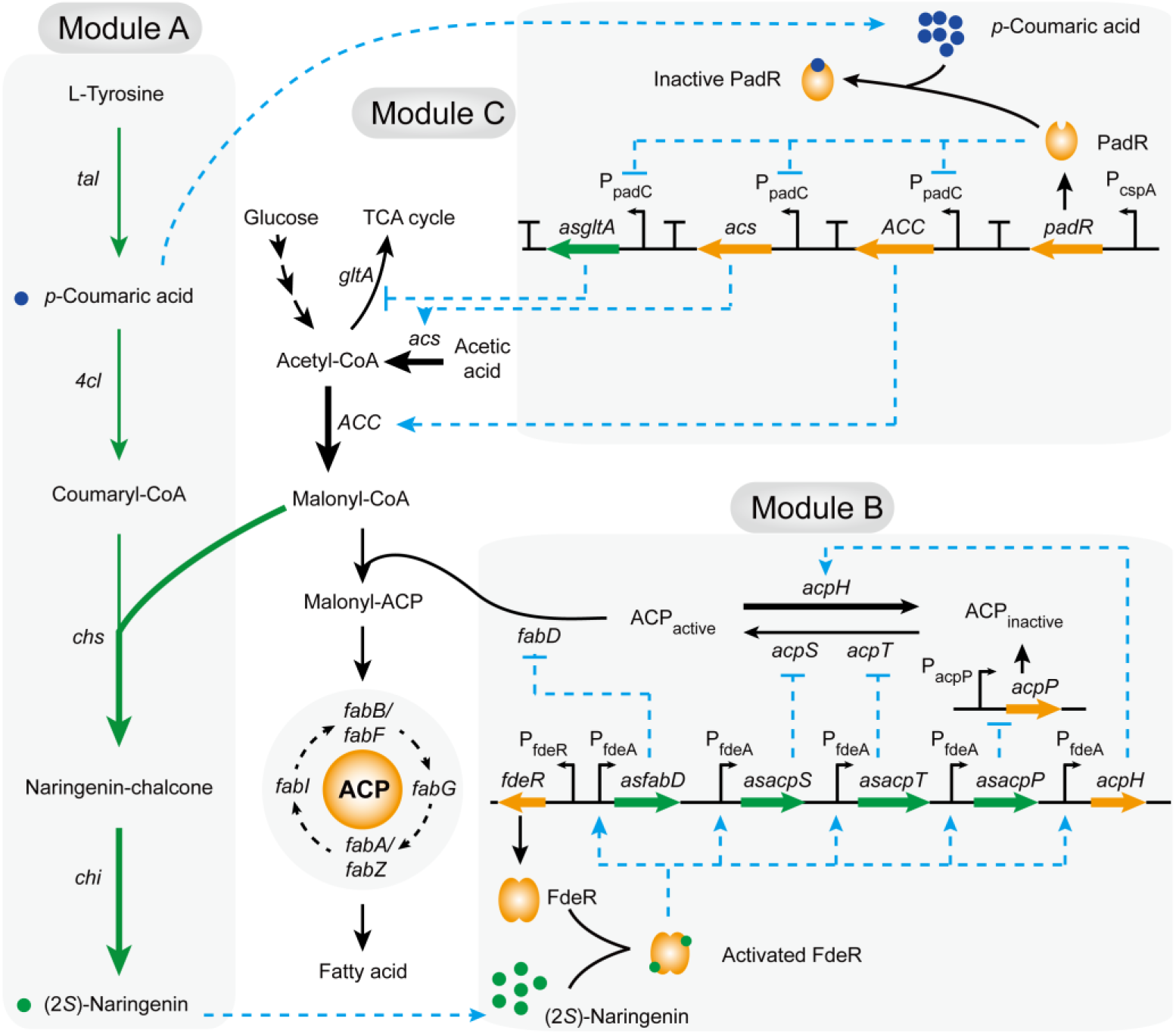
Schematic of the mechanisms for the NCOMB DRN. Module A: Heterologous (2*S*)-naringenin biosynthesis pathway; Module B: Fatty acid repression module. Module A and B could constitute to be a (2*S*)-naringenin-responsive amplification system to gradually elevate the repression level in malonyl-CoA consumption pathway over the course of the fermentation; Module C: Malonyl-CoA enhancing module.*p*-Coumaric acid-responsive pathway can dynamically enhance the supply of malonyl-CoA at stationary growth phase. Gene and protein annotations: tyrosine ammonia-lyase (*tal*), 4-coumarate: CoA ligase (*4cl*), chalcone synthase (*chs*), chalcone isomerase (*chi*), citrate synthase (*gltA*), acetyl-CoA synthetase (*acs*), acetyl-CoA carboxylase (ACC), acyl carrier protein (ACP), flavonoids-responsive transcriptional activator from *Herbaspirillum seropedicae* SmR1 (FdeR),*p*-coumaric acid-responsive transcriptional repressor from *Bacillus subtilis* (PadR), acyl carrier protein phosphodiesterase (*acpH*). *asfabD*, *asacpS, asacpT, asacpP*, and *asgltA* are anti-sense RNAs. Green arrows: constitutive expression pathway. Blue dash arrows: activation reactions; Blue dash blunt arrows: repression reactions. Bold arrows: enhanced metabolic fluxes.

## 2 Results

### 2.1 Construction and optimization of critical DRN biosensors

The (2*S*)-naringenin (FdeR based)-(*19*, *20*) and *p*-coumaric acid (PadR based)-(*21*) responsive biosensors are the central regulatory elements of our NCOMB DRN. To fine-tune the transcription of downstream target genes, it was necessary to engineer a moderate responsiveness and tight control of these elements. Although the responsive promoter architecture of P_fdeA_ (*19*) and P_padC_ (*22*, *23*) have been reported, information of the coresponding promoter strength is still unclear. Given that the length of 5’-UTR region for the P_fdeA_ and P_padC_ could change the secdonary structure of anti-senseRNA and affect the expression of target genes in our DRN design (**Fig. 1**), we first generated a series of truncated promoters to thoroughly investigate their impacts on the expression of corresponding genes of interest.

P_fdeA_ is a bidirectional promoter 288 bp in length and contains a dozen FdeR binding sites (T-N11-A) spread over a wide region (*19*). To dissect function of this element, we truncated P_fdeA_ in both directions based on the position of FdeR binding sites and generated 8 variants of P_fdeA_ (M-N) promoter actuated biosensors (**Fig. 2a and Supplementary Fig. 1a**). As a result, we identified that the shorter version of P_fdeA_(223-135), only 88 bp in length, exhibited around 4-fold higher signal-to-noise ratio than that of the orginal P_fdeA_ (288-1) when induced by 50 mg/L of (2*S*)-naringenin (**Fig. 2b and c**). Furthermore, the sensor with the truncated promoter P_fdeA_ (223-135) displayed a higher sensitivity over other constructs (**Supplementary Fig. 1b**). These properities can enable the newly designed (2*S*)-naringenin-responsive biosensor to express anti-sense RNAs and proteins (**Fig. 1**) at a very low level of (2*S*)-naringenin.

**Figure 2.**
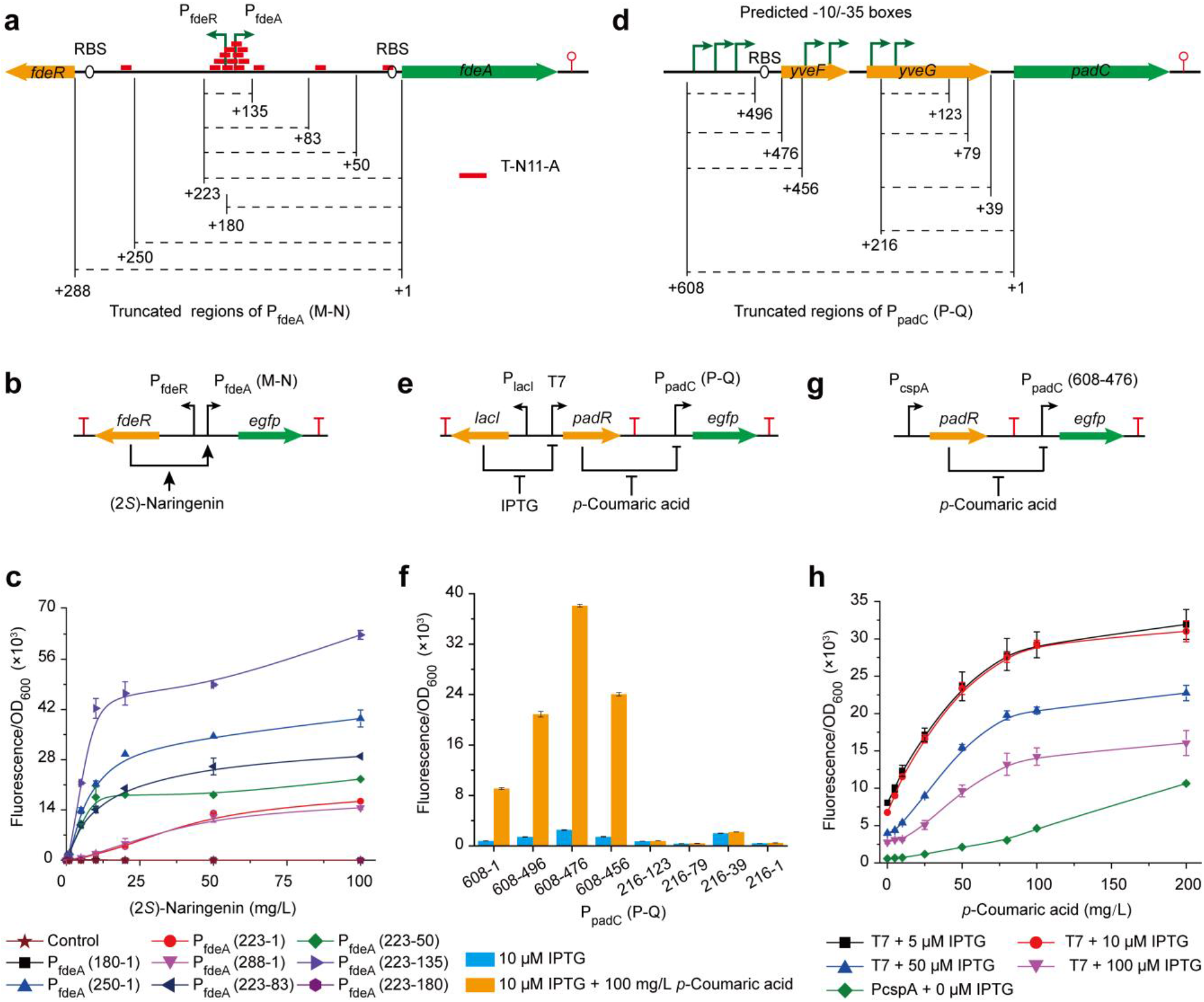
The structures and properties of (2*S*)-naringenin- and *p*-coumaric acid-responsive biosensors. **a**: The original structure of P_fdeA_ bidirectional promoter and truncated P_fdeA_ (M-N) regions. Red bar represents FdeR binding site (T-N11-A). **b**: The structure of (2*S*)-naringenin-responsive biosensor. **c**: The expression strength of promoter P_fdeA_ (M-N) when induced by various concentrations of (2*S*)-naringenin. Control represents strain only harboring pRSM-FdeR. Strains labeled with M-N represent constructs carrying pCDM-P_fdeA_(M-N)-GFP and pRSM-FdeR (**Supplementary Fig. 1a**). **d**: The predicted −35/−10 boxes of P_padC_ and truncated P_padC_(P-Q) regions. **e**: The structure of IPTG- and *p*-coumaric acid responsive biosensor. **f**: The expression strength of promoter P_padC_ (P-Q). **g**: The structure of *p*-coumaric acid-responsive biosensor. **h**: The properties of IPTG and/or *p*-coumaric acid inducible biosensors. Error bars represent standard error of biological triplicates.

Turning to the second promoter system, P_padC_, used to control the expression of *yveF*, *yveG*, and *padC* in an operon form, we predicted that the position of potential −35/−10 boxes by BDGP neural network promoter prediction software v2.2 (*24*) and idenfied 7 candidates sites (**Fig. 2d**). In a similar fashion with P_fdeA_, we established a promoter truncation series for P_padC_ based on the position of predicted −35/−10 boxes. Using the truncated promoter P_padC_(P-Q), 8 variants of *p*-coumaric acid and IPTG-responsive biosensors were constructed and evaluated (**Fig. 2e and Supplementary Fig. 1c**). As shown in **Fig. 2f**, the position of 608-476 was identified as a suitable region for protein expression, which demonstrated a 4.2-fold higher expression than that of the original P_padC_ (608-1). Furthermore, the shortest version (608-496) without the RBS exhibited an ideal expression strength, 2.3-fold of that of the original P_padC_ (608-1), which could be also used for expression of anti-sense RNA. To avoid the leaky expression and expensive inducer inherent in T7-based inducible control, we next employed a exponential phase-induced promoter P_cspA_ (*25*) to control the expression of PadR (**Fig. 2g**). Compared with the T7 promoter, P_cspA_ exbibited lower background with only 7.2% of that when 5 μM IPTG was added to the medium (**Fig. 2h**). Despite the lower total responsive level of P_cspA_ (only 33.3% the level of 5 μM IPTG-induced T7 controled biosensor (**Fig. 2h**)), the dynamic range of *p*-coumaric acid response was 18.2-fold higher in this system compared with the T7 controlled biosensor system. In addition, previous study has demonstrated that the expression level of P_cspA_ in exponential phase was 11.9-fold higher than in stationary phase (*25*). Taken together, these results highlight the use of P_cspA_ as a tighltly controlled sensor for *p*-coumaric acid accumulated in stationary phase.

### 2.2 Optimization of the fatty acid synthetic pathway

Most cellular malonyl-CoA in wild-type cells is consumed by the fatty acid synthetic pathway which requires an ACP to convert malonyl-CoA to fatty acyl-ACP for extending the saturated fatty acyl chain in each elongation cycles (*26*). Considering the important role of cofactor protein ACP in fatty acid synthetic pathway, the consumption of malonyl-CoA could be regulated by fine-tuning the activity of ACP. To do so, we overexpressed acyl carrier protein phosphodiesterase (*acpH*) to accelerate the transition from ACP_active_ to ACP_inactive_.(*27*) Furthermore, the anti-sense RNA of *acpS*, *acpT*, *acpP*, and *fabD* were also overexpressed to repress the translation of ACP, the transition from ACP_inactive_ to ACP_active_, and the formation of malonyl-ACP (**Fig. 1**). Through the combinatorial regulation of these genes, the concentration of malonyl-CoA was improved to at least 3.61-fold of that of control (**Fig. 3**). Through this analysis, we observed that higher malonyl-CoA concentrations led to lower biomass formation (**Fig. 3**). As a result, a proper dynamic level of *acpH* as *acpS*, and as *acpT* expression is necessary to balance between cell growth and malonyl-CoA accumulation.

**Figure 3.**
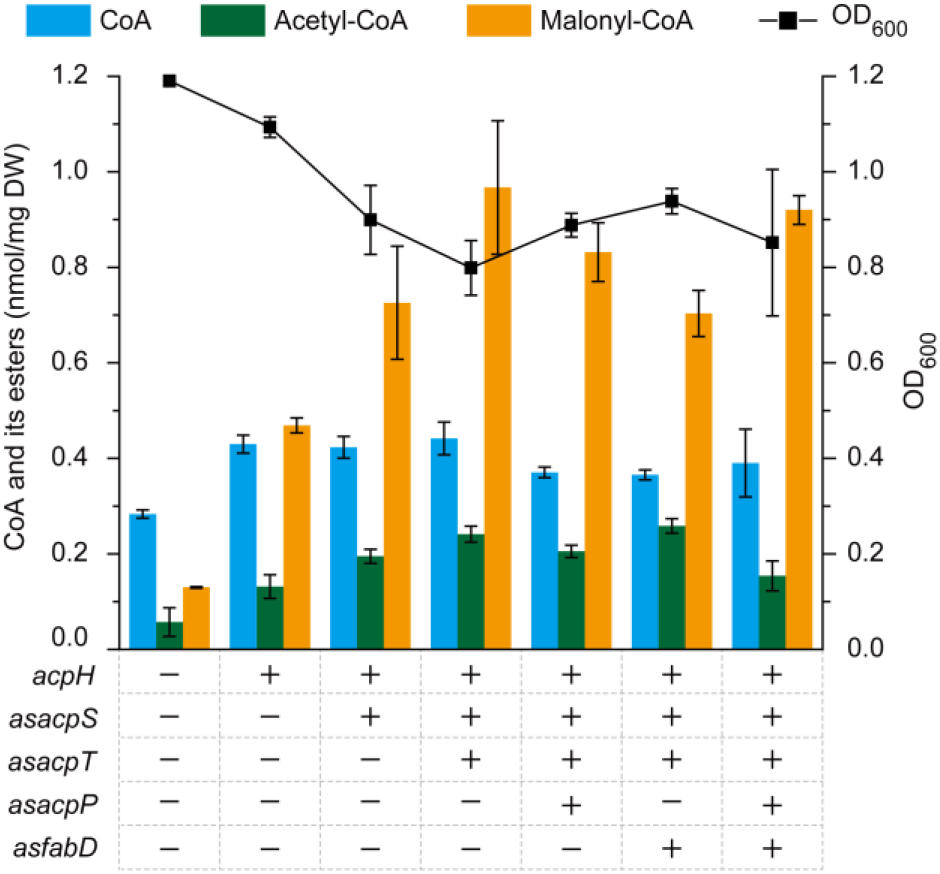
Fatty acid pathway optimization by (2*S*)-naringenin-responsive biosensor. Genes were overexpressed in pACM-FdeR plasmid under the control of P_fdeA_ (223-135) with 100 mg/L (2*S*)-naringenin induction. + or - represent the related genes were overexpressed or not, respectively. The control strain only harboring pACM-FdeR. Error bars represent standard error of biological triplicates.

Taken together, these results demonstrated that the repression of the transit form ACP_inactive_ to ACP_active_ state is more critical than repressing ACP translation and malonyl-ACP formation (**Fig. 1 and Fig. 3**). This could be explained by the rapid turnover of the AcpS and AcpT catalyzed reaction, thus resulting in almost exclusive ACP_active_ form (*28*). Furthermore, overexpression of AcpH not only suppresses the activity of ACP, but could also hydrolyze the 4’-phosphopantetheine moiety of ACP resulting in increasing amounts of CoA and acetyl-CoA after ACP repression (**Supplementary Fig. 2**) (*29*).

### 2.3 Designing a (2*S*)-naringenin-responsive amplifier to balance growth and malonyl-CoA accumulation

In order to dynamically balance cell growth and malonyl-CoA levels, we attempted to construct a (2S)-naringenin-responsive amplifier through coupling the previously optimized (2*S*)-naringenin synthetic pathway (Module A) (*14*) with the abovementioned fatty acid repression module (Module B) (**Fig. 4a**). This amplifier could dynamically repress the activity of ACP based on the cellular concentration of (2S)-naringenin. At early growth stages, the slight levels of (2S)-naringenin synthesized by module A will weakly activate module B and thus slowly repress the activity of ACP to produce just enough fatty acid levels to support biomass formation. During cell growth, increasing amounts of (2S)-naringenin will gradually enhance the ACP repression thus further increasing malonyl-CoA levels and (2S)-naringenin productivity. Thus, module B in our scheme is progressively activated to enable high metabolic flux toward (2S)-naringenin (**Fig. 4a**).

**Figure 4.**
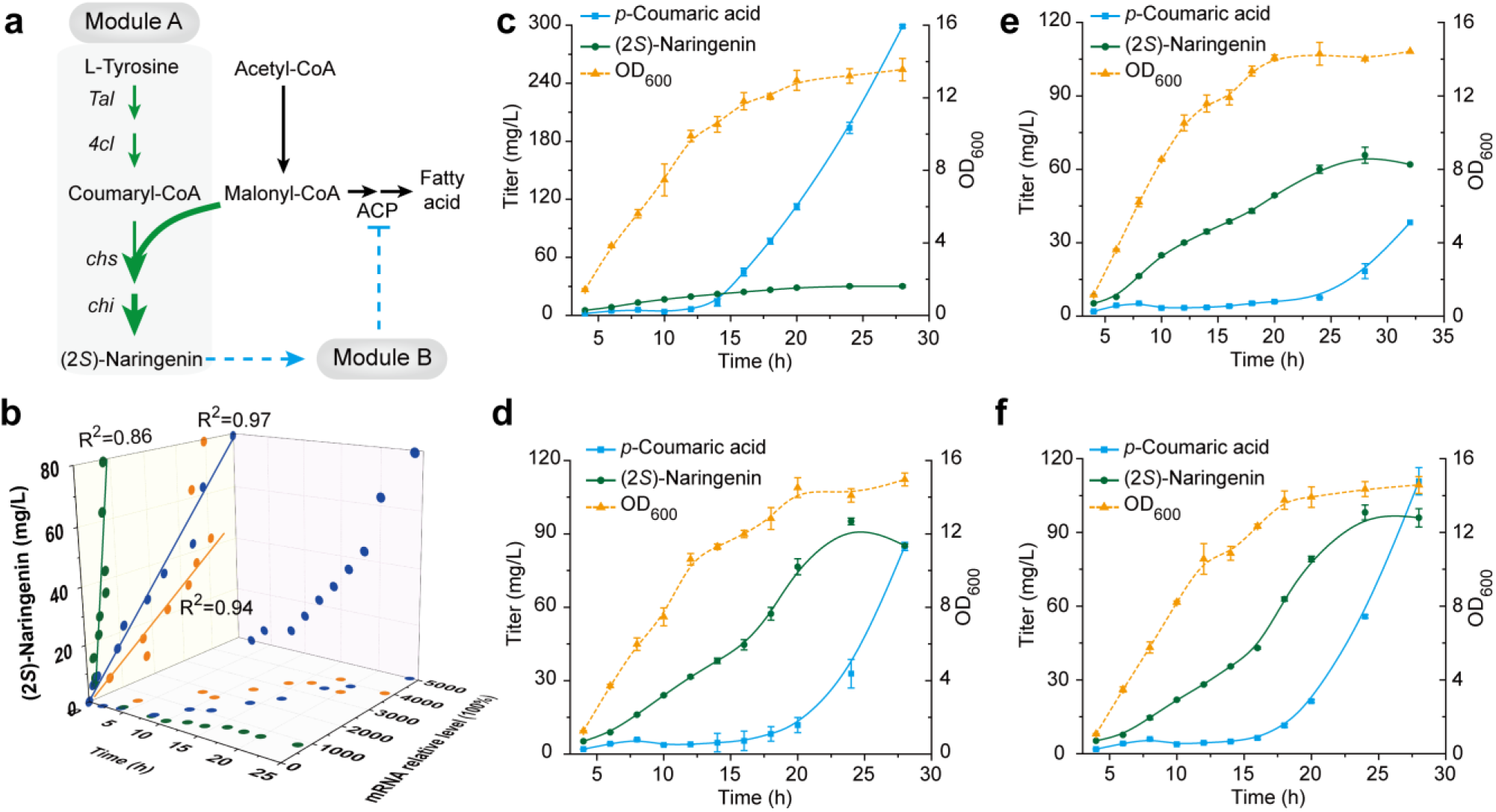
The mechanisms and properties of (2*S*)-naringenin-responsive amplifier. **a**: The mechanisms of (2*S*)-naringenin-responsive amplifier. **b**: The relative mRNA level and (2*S*)-naringenin titer of module A and B combined strain at different fermentation time. pACM-FdeR-*acpH*-as *acpT*-as *acpS* was used as module B. Blue, green, and orange points represent the genes of as *acpT*, as *acpS*, and *acpH*, respectively. **c-f**: The fermentation of different module A and B combined strains. The control strain (Nar_modA_) only harbors module A (**c**). Module A combined with pACM-FdeR-*acpH*-as *acpT*-as *acpS* (low copy number), pCOM-FdeR-*acpH*-as *acpT*-as *acpS* (medium copy number), pRSM-FdeR-*acpH*-as *acpT*-as *acpS* (high copy number) to generate strains of Nar_modAB-L_(**d**), Nar_modAB-M_ (**e**), and Nar_modAB-H_ (**f**), respectively. Error bars represent standard error of biological triplicates

To verify this concept, (2*S*)-naringenin titer and the expression level of *acpH*, as *acpS*, and as *acpT* was determined over the course of fermentation for strains with module A and B. Specifically, we observed a strong positive correlation between fermentation time, (2*S*)-naringenin concentration and the expression level of *acpH*, as *acpS*, and as *acpT* (**Fig. 4b**). The highest normalized expression levels of *acpH*, as *acpS*, and as *acpT* were 3945, 803, and 4983, respectively, when (2S)-naringenin titer reached 79.2 mg/L (**Fig. 4b**). These results indicate that the constructed amplifier can respond to a broad range of (2*S*)-naringenin concentration.

To further optimize cell growth, we utilized a three plasmid system with varying copy numbers to balance module B to enable proper fatty acid biosynthesis and cell growth. In doing so, the (2*S*)-naringenin titer was improved 2.2- to 3.2-fold compared with the control strain only expressing module A (Nar_modA_) without any impact on cell growth (**Fig. 4c-f**). As a result, the low copy number of module B was selected as the optimal module for (2*S*)-naringenin production. Comparing with the Nar_modA_, module B-combined strains significantly decreased the accumulation of intermediate *p*-coumaric acid after 14 h fermentation (**Fig. 4c-f**). This phenomenon could be the result of module B function which improved the supply of malonyl-CoA and thus increased the metabolic flux from *p*-coumaric acid-CoA to (2*S*)-naringenin. However, the *p*-coumaric acid of module B-combined strains rapidly accumulated and stop producing (2*S*)-naringenin after cell growth entered stationary phase (**Fig. d-f**). Therefore, the critical challenge left to further improve (2*S*)-naringenin production would be to efficiently utilize the accumulated *p*-coumaric acid in stationary phase.

### 2.4 Malonyl-CoA synthetic pathway optimization via *p*-coumaric acid-responsive biosensor

Previous study has demonstrated that the concentrations of acetyl-CoA and malonyl-CoA reached the highest levels in exponential phase and rapidly fell to 6.7% and 12.5% of the maximum level when entering stationary phase, respectively (*16*). Hence, we hypothesize that the accumulation of *p*-coumaric acid and decrease in (2*S*)-naringenin productivity observed in stationary phase was due to the limited availability of malonyl-CoA resulted from the reduced metabolic flux to biosynthesis of acetyl-CoA and malonyl-CoA in the stationary phase. To verify this hypothesis, we measured the concentration of malonyl-CoA of wild type *E. coli* BL21 at different growth phases and confirmed that malonyl-CoA reached its highest level in early growth stage and was almost undetectable in stationary phase (**Fig. 5a**). To enhance the intracellular level of malonyl-CoA, we constructed a dynamically regulated strain, Nar_modAC_ (**Fig. 5b**), through combining the *p*-coumaric acid-responsive dynamic regulation module (module C in **Fig. 1**) with module A. In doing so, the biosynthesis of malonyl-CoA would be enhanced by the overexpression of *acs*, *ACC*, and the anti-sense RNA of *gltA* (as *gltA*) when *p*-coumaric acid accumulated. In order to optimize this module C, the combinatorial overexpression of *acs*, *ACC*, and as *gltA* was evaluated and we found that overexpression of all genes facilitated the biosynthesis of (2*S*)-naringenin. Among all the constructs, simultaneous overexpression of *acs* and *ACC* leads to the highest (2*S*)-naringenin titer which was 1.5-fold higher than that of Nar_modA_ (**Fig. 5c**).

**Figure 5.**
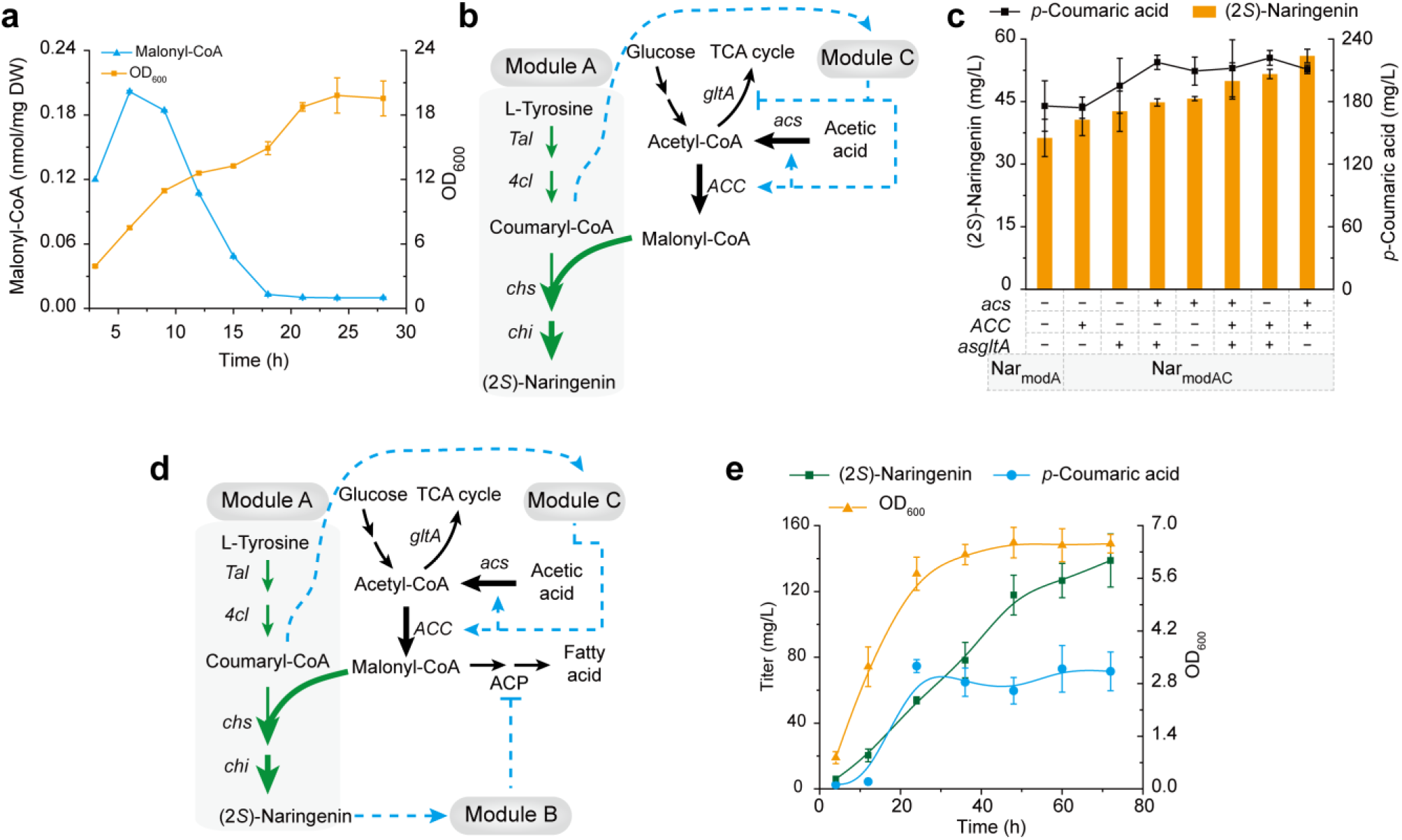
The mechanisms and properties of *p*-coumaric acid-responsive dynamic regulation strains. **a**: The malonyl-CoA concentration of wild type *E. coli* BL21 in different growth stages. **b**: The regulation mechanisms of strain Nar_modAC_. **c**: Optimization of module C through the combinatorial gene overexpression to generate various Nar_modAC_ constructs. Nar_modA_ was used as control. + or - represent the related genes were overexpressed or not, respectively. **d**: The regulation mechanisms of strain Nar_modABC_. Module B and C were pACM-FdeR-*acpH*-as *acpT*-as *acpS* and pRSM-PadR-*acs*-*ACC*, respectively. **e**: The (2*S*)-naringenin,*p*-coumaric acid, and OD_600_ level of Nar_modABC_ were determined. Error bars represent standard error of biological triplicates.

### 2.5 Complete DRN assembly for (2*S*)-naringenin production

Based on these collective results, and all modules in place, we combined the optimized modules A, B, and C to construct an artificial DRN which could simultaneously respond with (2*S*)-naringenin and *p*-coumaric acid to enhance the metabolic flux of consumption and biosynthesis of malonyl-CoA, respectively (**Fig. 1 and Fig. 5d**). Our system showed the steady production of *p*-coumaric acid and increasing level of (2*S*)-naringenin in the engineered strain Nar_modABC_ upon entry to stationary phase (**Fig. 5e**). In doing so, the final (2*S*)-naringenin titer reached 4.6-fold (138.9 mg/L) higher than that of Nar_modA_ (30.2 mg/L) and the titer of *p*-coumaric acid stayed nearly constant at around 70 mg/L. These results showcase that module C can effectively utilize *p*-coumaric acid to enhance the biosynthesis of (2*S*)-naringenin. However, the maximum OD_600_ of Nar_mod_ABC (6.5) was only 48.1% of that of Nar_modAB_ (13.5) suggesting that the combination of these three modules without fully expression optimization can inhibit the cell growth. Thus, this prompted us to apply a directed evolution strategy to fine-tune the expression level for DRN to rapidly and accurately balance the multiple artificial metabolic systems, and achieve the highest (2*S*)-naringenin titer without impaired cell growth.

### 2.6 Directed evolution to fine-tune DRN and achieve high level production

To tune the constructed DRN via directed evolution, we established a fluorescent-activated cell sorting (FACS)-based high-throughput screening approach (**Supplementary Fig. 3a**). A TtgR-based (2*S*)-naringenin biosensor (*30*) was constructed for FACS screening (**Fig. 6a**) leveraging the detection limit of this sensor (~ 220 mg/L) comparing with other biosensors, FdeR (10 mg/L; **Fig. 2b**) (*19, 20*) and (2S)-naringenin riboswitch (< 100 mg/L) (*31*, *32*). To normalize across cell size and general expression issues, we utilized a constitutively expressing far-red fluorescent protein mKate2 (*33*) (**Fig. 6a**). As shown in **Fig. 6b**, the TtgR biosensor exhibited a linear correlation between (2*S*)-naringenin concentration and GFP fluorescence when (2*S*)-naringenin was lower than 220 mg/L with a maximum induction of 16.6-fold. Past this concentration, GFP fluorescence decreased and we observed a negative correlation between the FRFP fluorescence and (2*S*)-naringenin concentration likely due to the cell burden of very high expression of GFP. As a result, we sought to identify cells that contain high green fluorescence and low red fluorescence as they are likely candidates for high titer.

**Figure 6.**
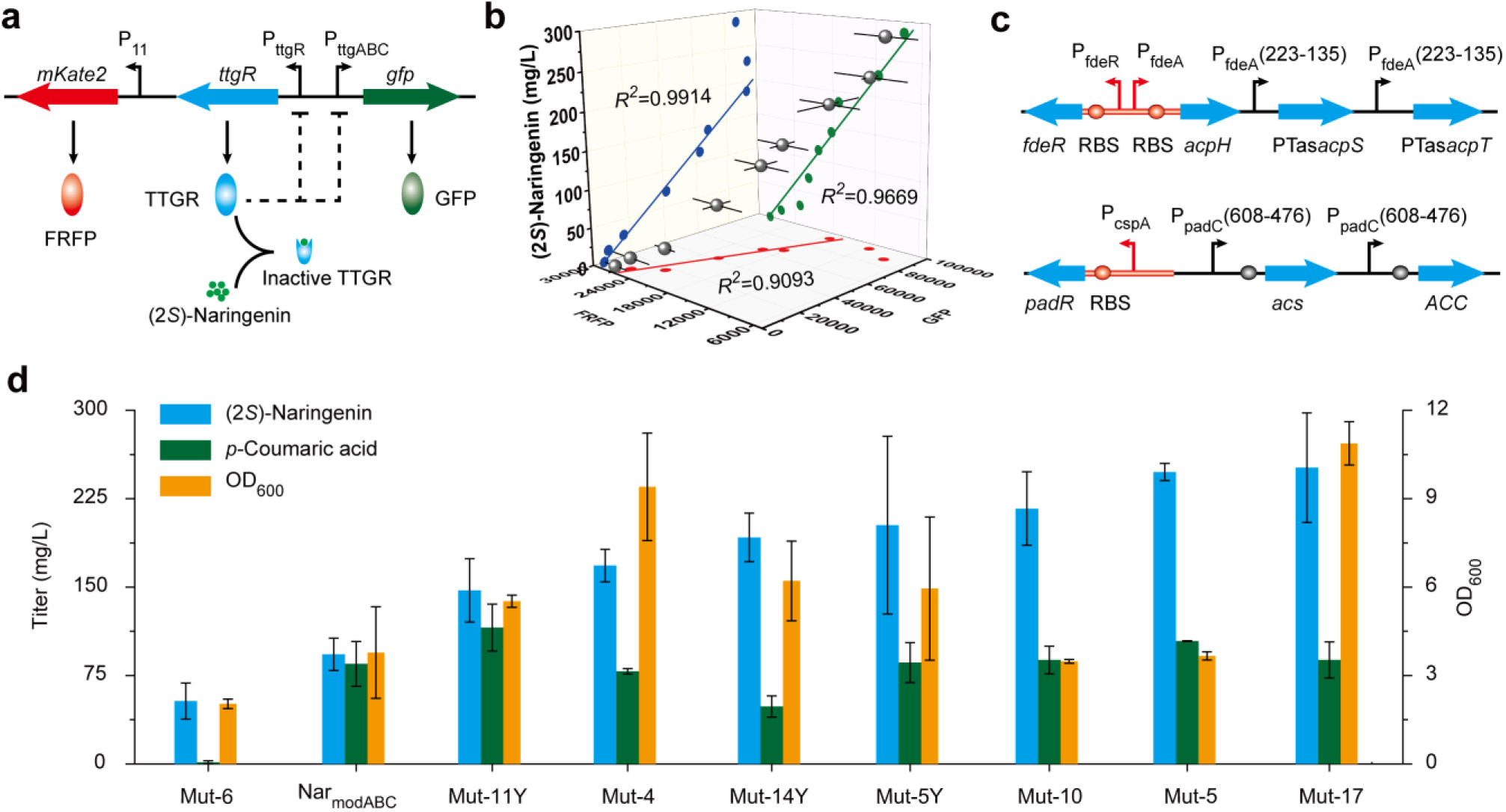
The FACS-based directed evolution. **a**: The structure of TtgR based (2*S*)-naringenin-responsive biosensor. **b**: The fluorescence levels of TtgR biosensor with different concentration of (2*S*)-naringenin inducer. **c**: The structure of module B and C. Red region represent the mutation area. **d**: The (2*S*)-naringenin,*p*-coumaric acid, and OD_600_ level of screened high titer strains. Error bars represent standard error of biological triplicates.

To fine-tune the function and performance of our DRN system, the P_fdeR_-P_fdeA_ of module B and P_cspA_ of module C were randomly mutated and then selected through FACS (**Fig. 6c**). The mutant library sizes were 2.02×10^7^ and 1.5×10^6^ for module B and C, respectively. After fermentation, the high naringenin-producing strains were sorted (**Supplementary Fig. 3a-c**) and then screened using a 96-well deep well plate assays (*14*). Among the variants tested, the best performing strain produced 251.4 mg/L (2*S*)-naringenin and 88.3 mg/L *p*-coumaric acid (**Fig. 6d**). When compared with the starting system containing Nar_modABC_, this mutant was improved 2.7-fold. Moreover, this strain was 1.6-fold improved in OD_600_. In order to characterize mutations in P_fdeR_-P_fdeA_ and P_cspA_ of derived mutants, the promoter sequences and their relative expression strength were further analyzed (more details in **Supplementary Notes**). Finally, we found that the high expression level of *acpH* is negative to (2*S*)-naringenin production.

## 3 Discussion

Dynamic control of metabolic pathways marked with precise control of enzyme expression levels as well as optimal cell growth has received attention in the last few years (*34*). In this study, we focus on optimizing and developing a NCOMB DRN to dynamically regulate the biosynthesis and conversion of malonyl-CoA in *E. coli* to direct it toward (2*S*)-naringenin production. In our three-part scheme, the constitutively expressed (2*S*)-naringenin synthetic pathway (module A) is coupled with a (2*S*)-naringenin-responsive biosensor (module B) and a stationary-phase malonyl-CoA enhancer (module C) to establish a dynamic amplifier balancing cell growth and malonyl-CoA consumption (**Fig. 1**). It should be noted that in our approach, ACP uniquely serves as the regulatory target instead of the commonly used fatty acid synthase (*35*). Like most synthetically designed circuits, it was necessary to optimize the interaction parameters in this system using FACS-based directed evolution. The underlying design and principles here have great potential to be extended to produce other valuable target compounds.

As a point of comparison with this work to others in the literature, it is well understood that intracellular malonyl-CoA content and impaired cell growth are the main challenges for (2S)-naringenin production (*35*). To address these limitations, previous studies have implemented a QS system to enhance malonyl-CoA production after sufficient cell growth. For example, Christina *et al.* established a bifunctional quorum-sensing DRN, which can dynamically enhance the expression level of TAL as well as 4CL and retard malonyl-CoA consumption rate (*34*). However, the reported (2*S*)-naringenin titer of best producer (125.9 mg/L) was only 50% of that of Mut-17 generated in this study (251.4 mg/L) (**Fig. 6d**). Comparison with NCOMB DRN, QS-based DRN suffered a signal distortion when transmission from cell density to gene expression. More specifically, we think the signal transition from cell density to malonyl-CoA concentration could not truly reflect the real demands for (2*S*)-naringenin production since cell growth state is also highly correlated to other factors including the energy and redox homeostasis (ATP/ADP and NADH/NAD+), the supply of carbon and nitrogen sources, pH, and temperature (*36*). In this study, the established DRN could avoid the distortion during signal transmission through directly sensing the accumulation level of (2*S*)-naringenin and *p*-coumaric acid to dynamically regulate gene expression, and finally achieve the goal of synchronizing cell growth and malonyl-CoA supplementation.

In this study, we successfully demonstrate that the optimized NCOMB DRN allows dynamic regulation of the metabolic fluxes in single cells according to their cell profiles. This strategy resulted in 8.4- and 1.6-fold improvement for (2*S*)-naringenin production and cell growth, respectively, compared to the control strain. We believe that the computational model (deep learning and machine learning modules (*37*, *38*))-based DRN with rational design and optimization can facilitate the improvement in the design accuracy and efficiency for more complex DRNs in the near future.

## 4 Materials and methods

### 4.1 Strains, medium, and culture conditions

*E. coli* JM109 and *E. coli* BL21 were used for plasmids construction and (2*S*)-naringenin production, respectively. LB medium was used to harvest cells for plasmid construction or fluorescence measurement. (2*S*)-naringenin fermentation was performed at 30°C with 220 rpm using MOPS minimal medium (supplemented with 10 g/L D-glucose, 1 g/L peptone, 3 mM L-tyrosine, and 4 g/L NH_4_Cl). Streptomycin (100 μg/mL), ampicillin (100 μg/mL), chloramphenicol (34 μg/mL), and kanamycin (50 μg/mL) were added to the media when needed. The strains, plasmids, and primers used in this study are listed in **Supplementary Table 1 and Table 2**.

### 4.2 Construction and characterization of biosensors

Restriction enzymes were purchased from Thermo Fisher Scientific (Waltham, MA). DNA ligase and PrimeSTAR HS DNA polymerase were purchased from Takara (Dalian, China). A ClonExpress II One Step Cloning Kit (Vazyme Biotech, Nanjing, China) or Gibson assembly method were used for plasmid construction. FdeR and P_fdeA_ promoter were synthesized and codon-optimized by Talen-bio Scientific (Shanghai) Co., Ltd.. Then, FdeR and P_fdeA_ promoter were amplified by primer pairs of FdeR(pACM)-F/FdeA-R, FdeR(pCOM)-F/FdeA-R, and FdeR(pRSM)-F/FdeA-R and cloned into *Apa*I/*Kpn*I, *Xba*I/*Kpn*I, and *Xba*I/*Kpn*I sites of pACM4, pCOM3, and pRSM3 to construct pACM-FdeR, pCOM-FdeR, and pRSM-FdeR, respectively. *gfp* was amplified by primer pairs of GFP-F/GFP-R and cloned into the *Spe*I/*Sal*I sites of plasmid pCDM4 to construct pCDM-GFP. Primer pairs of P_fdeA_(M)-F/P_fdeA_(N)-RBS-R or P_fdeA_(M)-F/P_fdeA_(1)-R were used to amplify different length of P_fdeA_ (M-N) promoters. Here, M and N represent the positions of amplified promoters, and the last 3’ base was defined as position 1. The amplified promoters were cloned into *Kpn*I/*Spe*I sites of pCDM-GFP to construct plasmids of pCDM-PfdeA(M-N)-GFP, *i.e.*, pCDM-PfdeA(180-1)-GFP, pCDM-PfdeA(223-1)-GFP, pCDM-PfdeA(250-1)-GFP, pCDM-PfdeA(288-1)-GFP, pCDM-PfdeA(220-50)-GFP, pCDM-PfdeA(220-83)-GFP, pCDM-PfdeA(220-135)-GFP, and pCDM-PfdeA(220-180)-GFP (**Supplementary Fig. 1a**). To characterize P_fdeA_ (M-N) promoters, the constructed pCDM-PfdeA(M-N)-GFP plasmids were transformed into control strain harboring a pRSM-FdeR plasmid (**Supplementary Fig. 1a**).

PadR and its terminator were amplified from the genome of *Bacillus subtilis* 168 by primer pairs of PadR-F/PadR-T-R. pRSM3 was digested by *Xba*I/*Kpn*I to obtain the 1.9 kb and 3.7 kb fragments which were subsequently ligated with PadR-terminator to construct plasmids of pRSM-PadR and pRSM-T7-PadR, respectively. An exponential phase expressed promoter P_cspA_ was amplified from the genome of *E. coli* MG1655 by primer pairs of cspA-F/cspA-R. Then, P_cspA_ was cloned into *Xba*I/*Asc*I site of pRSM-PadR to construct plasmid pRSM-P_cspA_-PadR. To simplify the structure of promoter P_padC_, primer pairs of PadC-P-F/PadC-Q-R were used to amplify different lengths of P_padC_ from the genome of *Bacillus subtilis* 168. Here, P and Q represent the positions of amplified promoters, and the last 3’-base was defined as position 1. The amplified promoters P_padC_ (P-Q) were ligated with *gfp* by fusion PCR, and then such promoters were cloned into *Bam*HI/*Spe*I sites of pRSM-T7-PadR to construct the plasmids of pRSM-T7-PadR-P_padC_(P-Q)-GFP (P-Q: 608-1, 608-496, 608-476, 608-456, 216-1, 216-39, 216-79, and 216-123). Furthermore, the P_padC_-(608-476)-*gfp* fragment was cloned into *Bam*HI/*Spe*I sites of pRSM-P_cspA_-PadR to construct the plasmids of pRSM-P_cspA_-PadR-P_padC_-(608-476)-GFP (**Supplementary Fig. 1c**).

To characterize the biosensors, strains were cultured in LB medium at 30°C. Various concentrations of (2*S*)-naringenin, IPTG, or/and *p*-coumaric acid were added to induce the expression of GFP when the OD_600_ reached 0.4-0.6. After 8 h induction at 30°C, cells were washed twice in PBS buffer (pH 7.4) and then fluorescence (488 nm excitation, 520 nm emission) and OD_600_ were measured by Cytation 3 imaging reader.

### 4.3 Pathway construction

Plasmid pUC57-asgltA that carried an anti-sense RNA gene of gltA (as *gltA*) was synthesized by Talen-bio Scientific (Shanghai) Co., Ltd.. Promoter P_fdeA_ (223-135) was amplified and cloned into *Not*I site of pUC57-asgltA to construct a pUC57-P_fdeA_-asgltA plasmid. Furthermore, complementary sequences of mRNA of *acpT*, *acpS*, *acpP*, and *fabD* were amplified from *E. coli* genome by primer pairs of asacpT-F/asacpT-R, asacpS-F/asacpS-R, asacpP-F/asacpP-R, and asfabD-F/asfabD-R, and subsequently cloned into *Nco*I/*Xho*I sites of pUC57-P_fdeA_-asgltA to construct plasmids of pUC57-P_fdeA_-asacpT, pUC57-P_fdeA_-asacpS, pUC57-P_fdeA_-asacpP, and pUC57-P_fdeA_-asfabD, respectively. Then, DNA fragments of P_fdeA_-asacpT, P_fdeA_-asacpS, P_fdeA_-asacpP, and P_fdeA_-asfabD were amplified from these plasmids by primer pairs of asRNA-F/asRNA-R. The *acpH* gene was amplified from the genome of *E. coli* MG1655 by primer pairs of acpH-F/acpH-R, and subsequently cloned into *Kpn*I/*Spe*I sites of pACM-FdeR, pCOM-FdeR, and pRSM-FdeR to construct plasmids of pACM-FdeR-acpH, pCOM-FdeR-acpH, and pRSM-FdeR-acpH, respectively. Finally, P_fdeA_-asacpT, P_fdeA_-asacpS, P_fdeA_-asacpP, and P_fdeA_-asfabD were digested by *Avr*II/*Sal*I, and subsequently cloned into *Nhe*I/*Sal*I sites of pACM-FdeR-acpH, pCOM-FdeR-acpH, and pRSM-FdeR-acpH by ePathBrick approach (*39*) to construct module B plasmids of pACM-FdeR-*acpH*-as *acpT*, pCOM-FdeR-*acpH*-as *acpT*, pRSM-FdeR-*acpH*-as *acpT*, pACM-FdeR-*acpH*-as *acpT*-as *acpS*, pCOM-FdeR-*acpH*-as *acpT*-as *acpS*, pRSM-FdeR-*acpH*-as *acpT*-as *acpS*, pACM-FdeR-*acpH*-as *acpT*-as *acpS*-as *acpP*, pACM-FdeR-*acpH*-as *acpT*-as *acpS*-as *fabD*, and pACM-FdeR-*acpH*-as *acpT*-as *acpS*-as *acpP*-as *fabD*. *E. coli* strains that harboring these plasmids were cultured in TB medium and induced by 100 mg/L (2*S*)-naringenin when OD_600_ reached 0.2. After 2 h induction, cells were harvested to measure the intracellular concentration of CoA and its esters.

Promoter P_padC_ (608-476) was amplified by primer pairs of padC-ACC-F/padC-476-ACC-R and padC-*Spe*I-F/padC476-acs-R, and subsequently ligated with ACC and acs to construct DNA fragments of P_padC_(608-476)-ACC and P_padC_(608-476)-acs, respectively. Promoter P_padC_ (608-496) was amplified by primer pairs of padC-*Spe*I-F/padC496-as *gltA*-R, and subsequently ligated with as *gltA* to construct DNA fragments of P_padC_(608-496)-asgltA. DNA fragments of P_padC_(608-496)-asgltA and P_padC_(608-476)-acs were cloned into *Xba*I/*Nde*I sites of pRSM-P_cspA_-PadR by ePathBrick approach (*39*) to construct plasmids of pRSM-PadR-acs, pRSM-PadR-asgltA, and pRSM-PadR-acs-asgltA. Then, P_padC_(608-476)-ACC was cloned into *Bam*HI/*Spe*I sites of pRSM-P_cspA_-PadR, pRSM-PadR-acs, pRSM-PadR-asgltA, pRSM-PadR-acs-asgltA to construct plasmids of pRSM-PadR-ACC, pRSM-PadR-acs-ACC, pRSM-PadR-asgltA-ACC, and pRSM-PadR-acs-asgltA-ACC, respectively.

### 4.4 FACS-based directed evolution

Far-red fluorescent protein (mKate2) and constitutive promoter P11 were amplified from plasmid pJKR-H-benM (*33*) with the primer pair of FRFP-F/FRFP-R. (2*S*)-Naringenin-responsive biosensor (including TtgR, GFP, and promoter P_ttgR_ and P_ttgABC_) was amplified from pJKR-H-ttgR (*30*) with the primer pair of TtgR-F/GFP-R. Then the PCR products of mKate2 and (2*S*)-naringenin biosensor were cloned into *Xba*I/*Bam*HI sites of pCDM-P_ssrA_-UTR_rpsT_-CHS-PUTR_glpD_-CHI by Gibson assembly method to construct the final plasmid pCDM-CHS-CHI-TtgR-GFP-FRFP.

GeneMorph II Random Mutagenesis Kit was used to generate mutation libraries of P_fdeR-mut_-P_fdeA-mut_ and P_cspA-mut_ with high mutation rate. Then the libraries were Gibson assembled with linearized plasmids of pACM-FdeR-acpH-asacpT-asacpS and RSM-PadR-acs-ACC to construct plasmid libraries of pACM-P_fdeR-mut_-FdeR-P_fdeA-mut_-acpH-asacpT-asacpS and pRSM-P_cspA-mut_-PadR-acs-ACC, respectively. Finally, these libraries were transformed into a *E. coli* strain which carried a constitutively expressing (2S)-naringenin synthetic pathway (module A) to generate a dynamic regulation library of *E. coli* Nar_lib_.

To perform the flow cytometry sorting, *E. coli* Nar_lib_ was cultured at 30°C and 220 rpm in MOPS medium for 24 h. Then cells were diluted to OD_600_ of 0.5. BD FACSAria III equipped with a 70 μm nozzle, 561 nm and 488 nm laser for excitation, and 610/20 nm and 530/30 nm filter was used for the sorting. The top 0.1% of single cells with high green fluorescence and low red fluorescence were sorted out from the mutant library (approximately 5×10^4^). The sorted cells were cultured in LB medium for 8 h at 37°C and then spread on agar plates. Approximately 3000 single colonies were picked into 96-deep-well plates containing 1 mL of MOPS medium in each well to screen high-titer strains by following our previously reported high-throughput screening method.(*14*) The high producers were further selected and fermented in 250 mL shaking flasks with biological triplicate. Finally, the mutation region of the selected high-titer strains was sequenced.

### 4.5 Real-time fluorescence quantitative PCR

Strains were cultured in MOPS medium at 30°C with 220 rpm orbital shaking. Cells were then harvested at 4°C with centrifuge speed 3000×*g* for3 min. mRNA was extracted from the harvested cells according to the manufacturer’s instructions of a RNAprep Pure Cell/Bacteria Kit (Tiangen, Beijing, China). PrimeScript RT reagent Kit with gDNA Eraser (TaKaRa, Japan) was used to reverse transcribe the extracted mRNA. SYBR Premix Ex Taq (Tli RNaseH Plus) was used for qRT-PCR on a LightCycler 480 II thermal cycler system (Roche, Mannheim, Germany). 16S rRNA gene was selected as internal standard.

### 4.6 Chromatography analysis methods

Equivalent volume of fermentation samples was mixed with ethanol at room temperature to extract (2*S*)-naringenin and *p*-coumaric acid. Then the mixtures were left at room temperature for 30 min to completely extract products. The supernatant of mixture was collected after centrifugation (4°C, 10000× *g*, 5 min) and filtered through 0.22 μm hydrophobic membranes. These supernatants were analyzed using an Agilent 1260 HPLC system. (2*S*)-Naringenin and *p*-coumaric acid were separated by a reverse-phase Gemini NX-C18 column (4.6 × 250 mm) at 25°C. Gradient elution method was the same as described in our previous report (*14*).

In order to measure the concentration of CoA and its thioester, strains were cultured in TB medium until OD_600_ reached 0.2, and then cells were harvested after induced by 100 mg/L of (2S)-naringenin for 2 h. CoA, acetyl-CoA, and malonyl-CoA were extracted according to the previously described method (*40*). Shim-pack VP-ODS (250 L ×2.0) UPLC column (Shimadzu, Tokyo, Japan) was used to separate CoA and its thioesters at 40oC with 15 mM ammonium formate in water (A) and 10 mM ammonium acetate in methanol (B) as mobile phases. The elution procedure was used as following: 0.2 mL/min flow rate, 10-25% B (vol/vol) for 5 min, 25-100% B (vol/vol) for 10 min, 100-10% B (vol/vol) for 1 min, maintaining 10% B (vol/vol) for 2 min. For quantitative analysis of CoA and its thioesters, the extracted ion chromatogram (EIC) peak areas were calculated by Shimadu LCMS-IT-TOF (Shimadzu, Kyoto, Japan). The detected EICs were: [M-H]^-^ = 766.5299 *m/z* for CoA, [M+H]^+^ = 810.5813 m/z for acetyl-CoA, [M-H]^-^ = 852.5764 *m/z* for malonyl-CoA. All the experiments were performed with biological triplicate. Means are reported, and the error bars represent standard deviation.

### 4.7 Nucleic acid sequences

The nucleic acid sequences of genes, promoters, and anti-sense RNAs were listed in **Supplementary Table 3** of Supplementary materials.

## Acknowledgements

This work was supported by the National Key R&D Program of China (2019YFA0905502), the National Science Fund for Excellent Young Scholars (21822806), the National Natural Science Foundation of China (31900066, 21877053, 31770097), the Air Force Office of Scientific Research under Award No. FA9550-14-1-0089, the National First-class Discipline Program of Light Industry Technology and Engineering (LITE2018-24), the Fundamental Research Funds for the Central Universities (JUSRP12056, JUSRP51705A).

## Author contributions

S. Z., J. Z., and Y. D. conceived the idea. S. Z. performed the experiments and data analysis. S. Z. and S.-F. Y. did the directed evolution. S.-F.Y. and P.H.N. did the 96-well plate screening. H.S.A. suggested the design idea of *p*-coumaric acid biosensor and supported the directed evolution. J. Z., Y. D., and H.S.A. critically reviewed the manuscript. S. Z., S.-F. Y., Y. D., and J. Z. wrote the paper. All authors reviewed, approved and contributed to the final version of the manuscript.

